# Pathological mitochondrial dysfunction mimics an aging pathway in budding yeast

**DOI:** 10.64898/2026.03.20.713207

**Authors:** Alexander Malyavko, Gilles Charvin

## Abstract

Mitochondrial dysfunction has long been associated with aging and linked to lifespan limitation across many species, including the budding yeast *Saccharomyces cerevisiae*. However, the widely used S288C laboratory background carries several polymorphisms that impair mitochondrial genome stability and function. Here, using a three-color reporter and single-cell microfluidics, we demonstrate how these mutations cause spontaneous transition to a state with severe mitochondrial deficiency characterized by low membrane potential, loss of heme biosynthesis, activation of iron regulon and morphological changes. Equally affecting young and old cells, this condition-dependent transition creates an apparent split in aging trajectories mimicking an age-dependent pathway. We further identify the mkt1-30D allele as a key genetic modifier of this pathological mitochondrial state. Together, these results suggest that mitochondrial dysfunction in this system reflects a genetic abnormality rather than an intrinsic aging process, calling for a reassessment of its role as a conserved hallmark. Our study also highlights how genetic defects can distort aging progression, potentially obscuring genuine age-associated phenotypes.

---

Aging is a ubiquitous feature of living organisms, yet its origins and exact nature still remains to be deciphered. Budding yeast provides a powerful model to identify and characterize the molecular mechanisms underlying cellular aging^1^. Because budding yeast divides asymmetrically, the mother cell can be distinguished from successive daughters over time, enabling quantitative single-cell analysis of aging trajectories. A mother cell typically undergoes ∼25 divisions, before entering senescence and eventually dying, a process referred to as replicative aging.

In an attempt to conceptualize aging, processes associated with its progression have been organized into twelve hallmarks^2^. Several of these hallmarks are considered to be conserved in replicatively aging yeast^3,4^. Perhaps, the most prominent example of such apparent conservation is mitochondrial dysfunction. Indeed, aging mother cells have been reported to exhibit loss of mitochondrial DNA (mtDNA) and mitochondrial membrane potential (MMP or ΔΨ)^5^, reduced heme biosynthesis^6^, activation of the iron regulon^5,7–9^, and mitochondrial fragmentation^6,10^. At the single-cell level, these changes have been associated with distinct aging trajectories: cells with compromised mitochondrial function produce small round daughters, whereas respiratory competent cells generate large elongated buds during several last divisions^1,6,11–13^.

However, most studies examining mitochondrial dysfunction during yeast aging have been performed in the S288C background, including the widely used BY4741 strain, which carries several polymorphisms that impair mitochondrial genome stability^14,15^. As a result, these strains frequently lose respiratory capacity even early in their replicative lifespan, leading to growth defects and formation of “petite” colonies. In addition, loss of MMP has been shown to occur stochastically and to be transmitted to daughters of aging mothers^16^. Together, these observations raise the possibility that the apparent divergence of aging trajectories may reflect the high rate of spontaneous petite formation in these strains, rather than an age-driven process.

In this study, we rigorously tested this idea by analyzing mitochondrial function in young and aged *S. cerevisiae* cells across several genetic backgrounds. First, we find that young dividing BY4741 cells are just as likely to develop severe mitochondrial dysfunction as cells in aging lineages. In contrast, only a small fraction of mothers exhibits strong loss of MMP in strains with stable mtDNA maintenance. We further uncover a crucial role of the MKT1 gene in determining the response to the loss of mitochondrial function. Finally, we show that the proportion of petite cells in the population influences the survival curve, providing a potential explanation for the “anti-aging” effects of some conditions. Altogether, our results demonstrate how genetic pathology can mimic an intrinsic aging process and suggest that mitochondrial dysfunction is not a hallmark of replicative aging in yeast.

## RESULTS

### The “mitotricolor” reporter strain for assessment of mitochondrial function during yeast aging

To capture distinct and complementary aspects of mitochondrial physiology, we engineered a BY4741 strain expressing three fluorescent reporters (hereafter mtBYtri, Fig. 1a). The NeonGreen protein fused to the mitochondrial targeting sequence of Cox15 (preCox15-NG), a mitochondrial inner membrane protein, reports on membrane potential, as protein import into mitochondrial matrix depends on MMP^17–19^. To monitor activation of the iron regulon, we expressed the red fluorescent protein ScarletI under the control of the promoter of *FIT2*, an iron-responsive gene induced upon mitochondrial dysfunction and during replicative aging^5,7,9^. Also, we used a nuclear-targeted infra-red protein (iRFP), as a proxy for heme levels^6^. iRFP fluorescence depends on the availability of biliverdin, a product of heme degradation, thereby linking its signal to cellular heme metabolism and mitochondrial function^6,20,21^.

**Figure 1.**
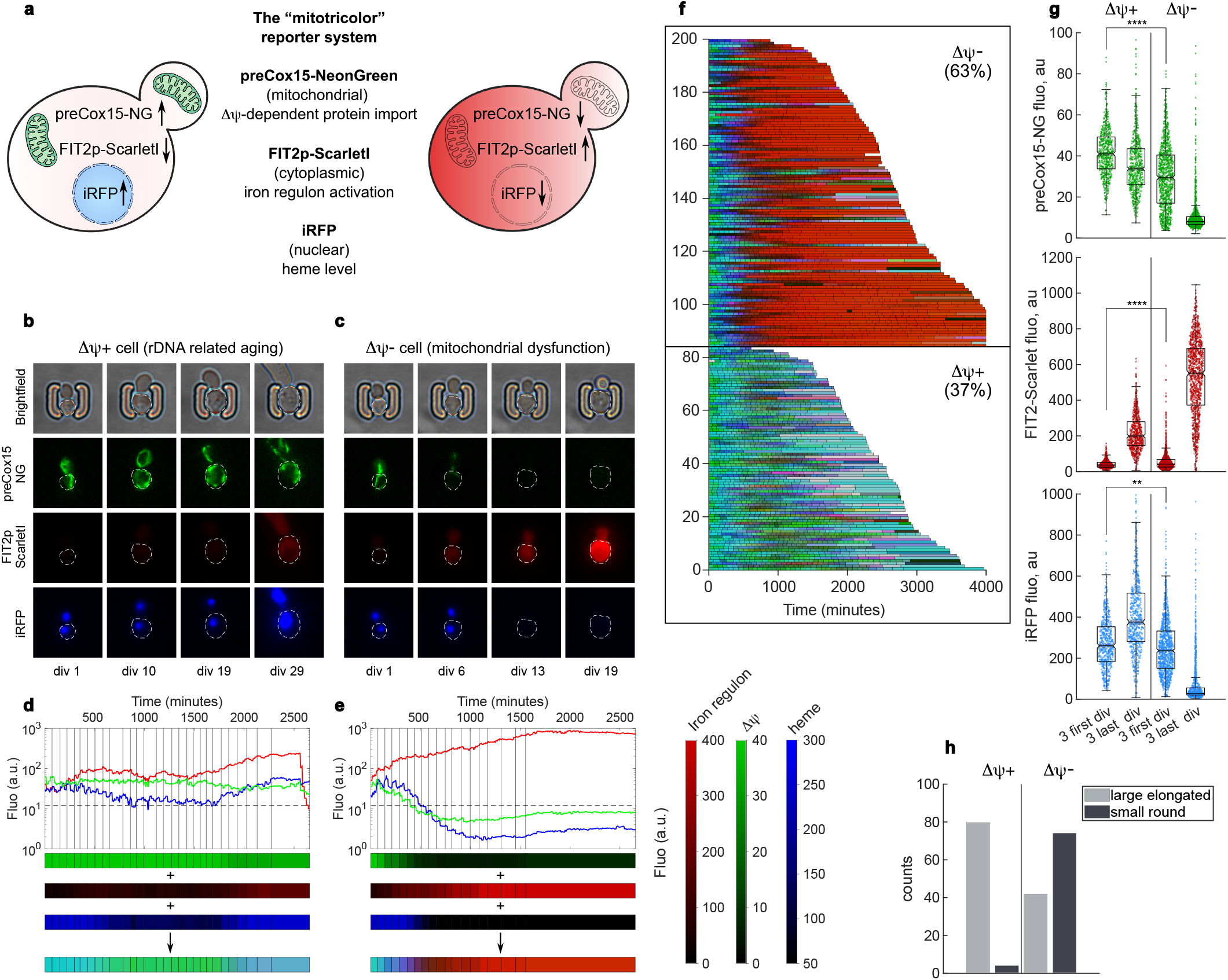
The “tricolor” reporter strain to detect mitochondrial dysfunction during yeast aging. **a**, Schematic representation of the principle behind the reporter system. **b**, Time-lapse sequence of brightfield and fluorescent images of aging mother cell with stable MMP (ΔΨ+ cell). White dashed lines contour mother cells; the number of divisions is indicated below. **c**, same as **b**, but the mother cell loses MMP during aging (ΔΨ-cell). **d**, Quantification of fluorescence of each marker in the ΔΨ+ mother cell depicted in **b**. Vertical black lines mark budding events. Bars below is another representation of the data: each rectangle is one cell cycle, color - mean fluorescence during that cycle. Three color bars can be concatenated into a single “RGB” bar representing the mitochondrial state of the cell. **e**, Same as **b**, but for the ΔΨ-mother cell. **f**, Single-cell trajectories (“RGB” bars from **d**,**e**) of the mtBYtri strain (200 representative trajectories randomly selected from three independent experiments) sorted by yeast lifespan (in minutes) and further sorted into two categories ΔΨ- or ΔΨ+ (numbers in parentheses indicate percentage of ΔΨ- or ΔΨ+ cells among 561 cells from three independent experiments). Cells were grown for ∼24 h in LOG phase before the aging experiment. **g**, Fluorescence (in a.u.) during three first and three last divisions (notches on boxplots delineate area (shaded in grey) of 95% confidence intervals around the median). ΔΨ+ n = 615, ΔΨ- n = 1068. ** - p <0.01, **** - p <0.0001 two-sample t-test. **h**, Number of mothers producing small round or large elongated buds among 200 mtBYtri cells in **f**.

To follow replicative aging at the single-cell level, we used a microfluidic device that enables continuous tracking of individual mother cells over time (“mother chip”)^22^. Using this approach, we observed that mtBYtri cells segregate into two distinct trajectories, consistent with previous reports^6,13,16^. A subset of cells maintained high preCox15-NG signal throughout their lifespan, showed only moderate activation of the iron regulon and exhibited a slight increase in iRFP fluorescence (ΔΨ+ cells, Fig. 1b,d). In contrast, another subset displayed severe mitochondrial dysfunction, characterized by near-complete loss of preCox15-NG signal, strong induction of FIT2p-ScarletI, and reduced iRFP fluorescence (ΔΨ-cells, Fig. 1c,e). These trajectories segregate clearly at the single-cell level (see ΔΨ- or ΔΨ+ cells on Fig. 1f), with coordinated changes across all three reporters (Fig. 1g, and Extended Data Fig. 1). Consistent with previous studies^6,11,13^, ΔΨ-mothers predominantly produced small round daughters during several terminal divisions, whereas ΔΨ+ mothers generated large elongated buds (Fig. 1h). Remarkably, many cells that adopt the ΔΨ-trajectory already exhibit reduced MMP and elevated FIT2p activity during the first divisions, suggesting that mitochondrial dysfunction is not triggered by age in these cells (Fig. 1g).

### Replicative age does not induce pathological mitochondrial dysfunction in yeast

Young BY cells can lose respiratory capacity with a certain probability^15^. Using our tricolor reporter system, we asked whether replicative age increases the probability of mitochondrial failure and/or exacerbates its phenotypic consequences once it occurs. To this end, we designed a microfluidic device (“daughter chip”) whose trap geometry makes it possible to follow either aging or rejuvenating cells depending on their budding pattern (Fig. 2a,b). In wild-type haploid BY4741 cells, budding is predominantly axial, so newly formed daughters emerge near the previous bud site and mother cells generally remain at the bottom of the trap, allowing aging lineages to be tracked over time (Fig. 2a). By contrast, *bud4Δ* cells adopt a bipolar budding pattern^23^ and are therefore frequently replaced by their daughters, such that the cells retained at the bottom of the trap belong to continuously rejuvenating lineages (Fig. 2b).

**Figure 2.**
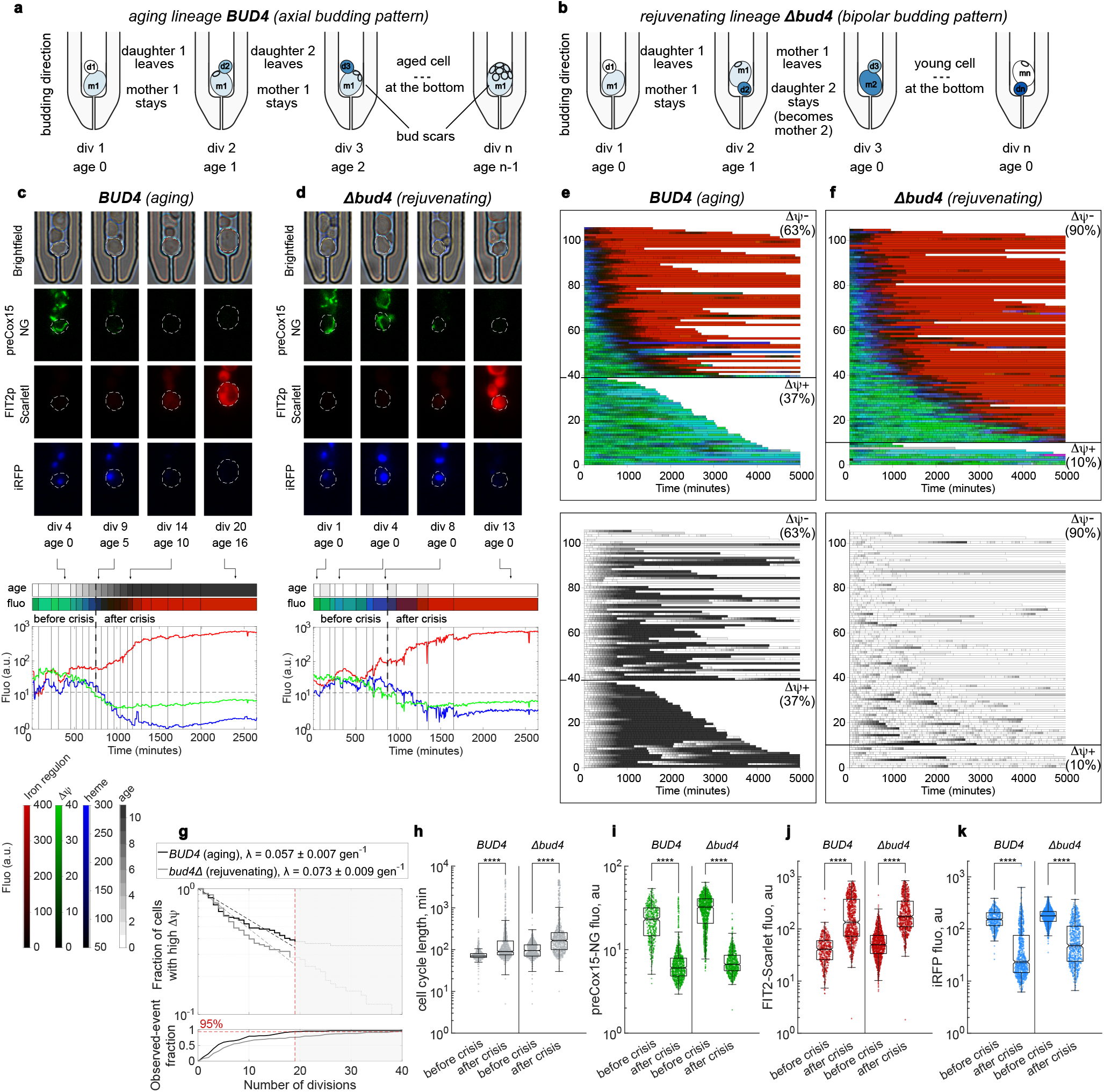
Similarities of mitochondrial dysfunction between aging and constantly rejuvenating lineages. **a-b**, Schematic of WT (**a**) and *bud4Δ* (**b**) cell divisions in the “daughter” microfluidics chip. Color represents the cell identity. “div” – division number, “age” – replicative age. WT predominantly buds in one direction and when captured in the configuration shown in the figure, the daughters it produces will leave the trap and the mother stays at the bottom. *bud4Δ* cells regularly switch the direction of budding and mother cell will be frequently replaced by its daughter (see how d2 replaces m1 in the example in **b**), resetting the replicative age. **c**, (top panel) Time-lapse sequence of brightfield and fluorescent images of a WT aging mother ΔΨ-lineage in the “daughter” chip. White dashed lines contour tracked cells at the bottom of the trap. Note several replacement events before the mother cells started budding towards the exit of the trap (leading to a gap between the divisions and the replicative age (“div 4” and “age 0”)). (bottom panel) Quantification of fluorescence of each marker in the ΔΨ-aging lineage. Vertical black lines mark budding events. Above is shown the “RGB” bar of the data (“fluo”) along with the bar with color-coded replicative age (“age”). Vertical dashed line marks the “crisis”. Below are color maps used in the figure. **d**, Same as **c**, but with a *bud4Δ* daughter ΔΨ-cell lineage. **e**, (top panel) Single-cell trajectories of the *BUD4* mtBYtri strain with fluorescence data from the experiment in the “daughter” chip sorted into two categories ΔΨ- or ΔΨ+ (numbers in parentheses indicate percentage of ΔΨ- or ΔΨ+ lineages among 106 lineages from the experiment). ΔΨ+ lineages were sorted by yeast lifespan (in minutes); ΔΨ-lineages were sorted by time of mitochondrial dysfunction onset (preCox15-NG signal < 12 a.u.). Cells were grown for ∼24 h in LOG phase before the aging experiment. (bottom panel) Single-cell trajectories of the *BUD4* mtBYtri strain with replicative age data. **f**, Same as **e**, but with the *bud4Δ* mtBYtri strain. 105 lineages were analyzed in the experiment. **g**, Time to ΔΨ loss in daughter-chip lineages. Kaplan-Meier estimates were computed by treating lineages that died or remained ΔΨ+ at the end of the experiment as right-censored observations. Quantitative fits were restricted to the common informative window (first 19 divisions), which contains 95% of observed events in *BUD4* daughter lineages. Solid lines show the Kaplan-Meier estimates within this window; dashed lines show exponential fits; dotted extensions indicate the censored late tail that was not used for fitting. The lower panel shows the cumulative fraction of observed ΔΨ-loss events used to define the cutoff. **h-k**, Log10-transformed cell cycle length (in minutes) and fluorescence (in a.u.) during divisions before and after mitochondrial dysfunction onset (“before or after crisis”) (notches on boxplots delineate area (shaded in grey) of 95% confidence intervals around the median). Only ΔΨ-lineages were analyzed. *BUD4* before crisis n = 478, *BUD4* after crisis n = 921, *bud4Δ* before crisis n = 1139, *bud4Δ* after crisis n = 751. **** - p <0.0001 two-sample t-test.

Consistent with the results obtained in the “mother chip”, many aging mtBYtri lineages in the “daughter chip” followed ΔΨ-trajectories (example in Fig. 2c). Remarkably, signs of mitochondrial dysfunction were also detected among rejuvenating lineages. In one representative lineage composed of cells with replicative age below 2, after ∼7 normal divisions daughter cells began to lose MMP and heme levels, activate iron regulon, and progressively slow down cell division until a complete arrest after the 13^th^ division (Fig. 2d).

Analysis of more than 100 lineages per genotype revealed that rejuvenating mtBYtri *bud4Δ* daughter lineages were just as prone to losing mitochondrial membrane potential as aging mtBYtri *BUD4* daughter lineages, despite remaining at low replicative age throughout the experiment (Fig. 2e,f and Extended Data Fig. 2a). The final fraction of ΔΨ-lineages was in fact higher in *bud4Δ* than in *BUD4* cells (∼90% versus ∼60%; Fig. 2e,f), an observation strongly suggesting that ΔΨ loss is not simply a hallmark of replicative aging. A parsimonious explanation for the lower final fraction of ΔΨ-lineages in *BUD4* is that an age-dependent competing hazard removes some lineages before ΔΨ loss can be observed, thereby reducing the apparent endpoint fraction without implying a lower intrinsic propensity to undergo ΔΨ loss. Consistent with this interpretation, mtBYtri *bud4Δ* cells followed as aging lineages in the “mother chip” produced fewer ΔΨ-lineages than rejuvenating mtBYtri *bud4Δ* cells in the “daughter chip” (Extended Data Fig. 2b versus Fig. 2f), indicating that endpoint fractions are strongly shaped by competing lineage loss.

To quantify the onset of mitochondrial dysfunction while accounting for censoring, we analyzed time to ΔΨ loss using Kaplan-Meier estimates, treating lineages that died or remained ΔΨ+ until the end of the experiment as right-censored observations. Because the late tails of the distributions are dominated by censored trajectories, we restricted quantitative analysis to a common informative window spanning the first 19 divisions in the “daughter chip” (and 16 in the “mother chip”; Extended Data Fig. 2c), which already contains 95% of observed events (Fig. 2g and Extended Data Fig. 2c). Within this interval, the data were consistent with an exponential survival model and were not better described by a Weibull model with a free shape parameter k. A Weibull model would capture a systematic increase (k > 1) or decrease (k < 1) in the probability of ΔΨ loss with division number if present. However, we did not detect a statistically significant change in the probability of ΔΨ loss with replicative age (*BUD4* daughter, k = 0.86, 95% CI, 0.70-1.07, p = 0.17; *bud4Δ* daughter, k = 1.01, 95% CI, 0.84-1.23, p = 0.90; Extended Data Fig. 2d). Exponential fits yielded rates of the same order in aging *BUD4* and rejuvenating *bud4Δ* daughter lineages (λ = 0.055 ± 0.007 gen^-1^ versus 0.073 ± 0.008 gen^-1^; Fig. 2g), indicating that ΔΨ loss arises with similar kinetics in rejuvenating and aging lineages, rather than being specifically accelerated by replicative aging.

In addition, we observed quantitatively similar cell cycle slowdown, MMP loss, iron regulon activation and reduction in heme levels following mitochondrial crisis in both aging and non-aging mtBYtri lineages (Fig. 2h-k). By contrast, an increase in cell size was observed only in mtBYtri *BUD4* lineages, suggesting that this feature is linked to replicative age independently of mitochondrial health (Extended Data Fig. 2e). Collectively, these results strongly suggest that severe mitochondrial dysfunction is neither induced nor promoted by replicative age in the BY4741 strain.

### Loss of mitochondrial function during aging in BY4741 is linked to its impaired mtDNA maintenance system

Strains of S288C genetic background, from which BY4741 is derived, harbour several polymorphisms detrimental to mtDNA replication and respiratory capacity (Extended Data Fig. 3a). Mutations in the *SAL1, CAT5* and *MIP1* genes are largely responsible for the high rate of petite formation^15^, while a variant in *MKT1* reduces the expression of many nuclear-encoded mitochondrial proteins^24^, and truncation of the heme-responsive transcription factor *HAP1* alters respiratory metabolism^25^.

To test whether these polymorphisms drive the emergence of ΔΨ-aging trajectories in BY4741, we analyzed mitochondrial states during replicative aging in strains carrying functional versions of the alleles. We selected the vineyard isolate RM11-1a, originally used for QTL analysis of the petite phenotype^15^, and the DHY213 strain^26^. The latter is essentially BY4741 in which five aforementioned genes have been restored (Extended Data Fig. 3a), along with two additional mutations (in *TAO3* and *RME1*) that improve sporulation efficiency. We observed a markedly reduced petite formation frequency in the tricolor versions of both strains (mtRMtri and mtDHYtri, Extended Data Fig. 3b). Strikingly, none of the aging lineages of these strains resembled ΔΨ-trajectories comparable to those observed in mtBYtri, characterized by the simultaneous loss of MMP and heme levels together with activation of the iron regulon (compare Fig. 3c,e to Fig. 3a). Instead, fluorescent reporter dynamics in the vast majority of mtRMtri and mtDHYtri mother cells showed only moderate changes, reminiscent of the ΔΨ+ trajectories of mtBYtri (compare Fig. 3c,e to Fig. 3a, see Fig.3g for the numerical data).

**Figure 3.**
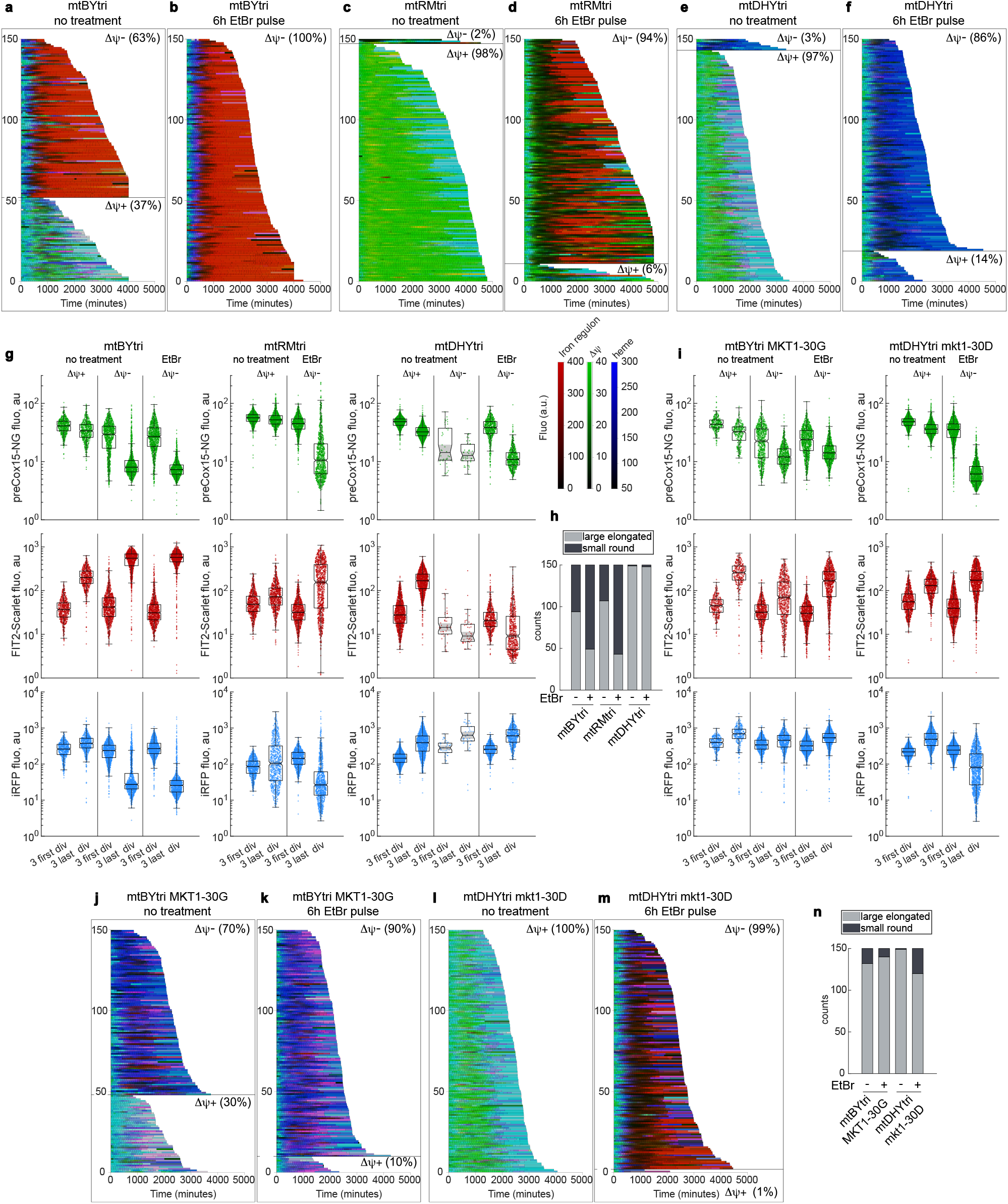
Mitochondrial dysfunction in the “petite-less” yeast strains. **a-f**, Single-cell trajectories (150 representative lineages randomly selected from one to three independent experiments) of the indicated strains with fluorescence data from experiments in the “mother” chip with or without EtBr pretreatment 6h prior to the loading onto chip. Lineages were sorted into ΔΨ- or ΔΨ+ (numbers in parentheses indicate percentage of ΔΨ- or ΔΨ+ lineages among n lineages. mtBYtri no treatment n = 561 (3 experiments), EtBr n = 358 (2 experiments); mtRMtri no treatment n = 270 (2 experiments), EtBr n = 359 (2 experiments); mtDHYtri no treatment n = 480 (2 experiments), EtBr n = 265 (2 experiments)) Cells were grown for ∼24 h in LOG phase before the aging experiment. **g**, Log10-transformed fluorescence (in a.u.) during three first and three last divisions (notches on boxplots delineate area (shaded in grey) of 95% confidence intervals around the median). mtBYtri no treatment ΔΨ+ n = 615, ΔΨ-n = 1068 (3 experiments), EtBr ΔΨ-n = 1071 (2 experiments); mtRMtri no treatment ΔΨ+ n = 798 (2 experiments), EtBr ΔΨ-n = 1008 (2 experiments); mtDHYtri no treatment ΔΨ+ n = 1397, ΔΨ-n = 42 (3 experiments), EtBr ΔΨ-n = 681 (2 experiments). Statistical analysis (two-sample t-test) of the data is in Supplementary Table 1. **h**, Number of mothers producing small round or large elongated buds among 150 trajectories shown in **a-f. i**, Log10-transformed fluorescence (in a.u.) during three first and three last divisions (notches on boxplots delineate area (shaded in grey) of 95% confidence intervals around the median). mtBYtri MKT1-30G no treatment ΔΨ+ n = 315, ΔΨ-n = 756 (2 experiments), EtBr ΔΨ-n = 1023 (2 experiments); mtDHYtri mkt1-30D no treatment ΔΨ+ n = 960 (2 experiments), EtBr ΔΨ-n = 1371 (3 experiments). **j-m**, Single-cell trajectories with fluorescence data (150 representative lineages randomly selected from two to three independent experiments) of the indicated strains from experiments in the “mother” chip. Lineages were sorted into ΔΨ- or ΔΨ+ (numbers in parentheses indicate percentage of ΔΨ- or ΔΨ+ lineages among n lineages. mtBYtri MKT1-30G no treatment n = 357 (2 experiments), EtBr n = 378 (2 experiments); mtDHYtri mkt1-30D no treatment n = 320 (2 experiments), EtBr n = 461 (3 experiments). Statistical analysis (two-sample t-test) of the data is in Supplementary Table 1. **h**, Number of mothers producing small round or large elongated buds among 150 trajectories shown in **j-m**.

However, even young mtRMtri and mtDHYtri ΔΨ+ mothers exhibited noticeable differences in fluorescence, resulting in a greener hue of tricolor bars compared to mtBYtri (Fig. 3a,c,e). To ensure that our system could reliably detect mitochondrial dysfunction in mtRMtri and mtDHYtri despite this altered baseline, we induced respiratory deficiency by pretreating cells with a short pulse of ethidium bromide (EtBr) prior to the aging experiment. This treatment causes extensive mtDNA mutagenesis rendering most cells petite (Extended Data Fig. 3c). Following EtBr treatment, 100% of mtBYtri aging lineages transitioned to the ΔΨ-state (Fig. 3b), confirming that corresponding trajectories observed in untreated strain result from mtDNA loss or mutation.

Interestingly, although respiratory-deficient mtRMtri and mtDHYtri lost preCox15-NG signal, the behavior of the other two reporters was markedly different from mtBYtri. In mtRMtri, iRFP fluorescence was lost, but FIT2p activation was much more attenuated than in mtBYtri lineages (Fig. 3b,d); some mtRMtri mother cells even regained preCox15-NG signal without strong iron regulon activation (Fig. 3d). In contrast, ΔΨ-mtDHYtri mothers did not activate FIT2p expression (even lost the residual activation of the ΔΨ+ untreated cells), while maintaining iRFP fluorescence (Fig. 3e,f). Based on these criteria, we were able to identify a small number of lineages among mtRMtri and mtDHYtri cells that likely lost respiratory capacity during normal aging (labeled ΔΨ-in Fig. 3c,e), but these represented only a small fraction of all the cells in the experiment. Thus, the apparent divergence of fate trajectories in BY4741 largely reflects its impaired mtDNA maintenance.

Pretreatment with EtBr increased the proportion of mtBYtri mother cells producing small round daughters, further strengthening the association between this morphological subtype and mitochondrial dysfunction (Fig. 3h). Surprisingly, a substantial fraction (∼30%) of ΔΨ+ mtRMtri cells also produced small round daughters (Fig. 3h), despite no obvious loss of mitochondrial function. We note, however, that small round buds are frequently observed throughout the lifespan of mtRMtri cells, even in mothers that eventually start producing large elongated daughters (Extended Data Fig. 3d). In contrast, aged mtDHYtri cells produced almost exclusively large elongated buds, with or without EtBr pretreatment (Fig. 3h). These observations suggest that the link between mitochondrial dysfunction and morphological phenotype during replicative aging yeast breaks down in strains with low rates of petite formation.

### A defective MKT1 allele is largely responsible for the uncontrolled response to the loss of mitochondrial function in BY4741

As described above, mtBYtri and mtDHYtri strains respond differently to EtBr-induced loss of MMP: In mtBYtri this is accompanied by strong activation of iron regulon, complete loss of iRFP fluorescence, and a pronounced decrease in MMP, whereas these responses are largely absent or attenuated in mtDHYtri (Fig. 3a,b,e-g). In addition, only mtBYtri ΔΨ-mothers produced small round daughters (Fig. 3h). Given the limited genetic variation between the two strains, we wondered whether specific allelic variants could account for these phenotypic differences. To address this, we analyzed responses to EtBr pulse in a set of random spores derived from mtBY/DHYtri crosses (Extended Data Fig. 4). This analysis yielded *SAL1* and *MKT1* as candidate loci associated with differential reporter responses (compare crosses #2 and #13 in Extended Data Fig. 4b,c).

Importantly, replacing only the dysfunctional mkt1-30D allele in the mtBYtri strain with the MKT1-30G allele from DHY213 (mtBYtri MKT1-30G) strongly reduced the iron regulon activation following EtBr treatment (Fig. 3b,k). In these cells, heme levels were maintained and the decrease in MMP was more moderate, closely resembling the mtDHYtri phenotype (Fig. 3e-k). We note, however, that MKT1-30G did not reduce the percentage of ΔΨ-lineages is the untreated sample (63% of ΔΨ-in mtBYtri vs 70% of ΔΨ-in mtBYtri MKT1-30G, Fig. 3a,j), suggesting that this allele does not decrease the rate of mitochondrial function loss, but rather modulates the phenotype of the ΔΨ-cells. Remarkably, the majority of old mtBYtri MKT1-30G mother cells produced large elongated daughters despite the loss of respiratory function (Fig. 3n).

Conversely, introduction of the dysfunctional mkt1-30D allele into the mtDHYtri strain caused its response to mtDNA loss to resemble that of wild-type mtBYtri. In EtBr-treated cells, this was characterized by reduced heme production, activation of the iron regulon and a more pronounced decrease in MMP (Fig. 3l,m and numerical data in Fig. 3i). Notably, a subset of the mtDHYtri mkt1-30D mothers (∼20%) produced small round daughters (Fig. 3n), a phenotype that was almost never observed in the parental mtDHYtri strain (Fig. 3h). However, the magnitude of FIT2p expression in old cells and the proportion of small round daughters remained lower than in mtBYtri, suggesting that sal1-1 (and possibly other polymorphisms) contributes to the mtBYtri ΔΨ-phenotype. Collectively, these results demonstrate a key role for *MKT1* in determining the severity of the response to loss of respiratory capacity in yeast, without affecting the frequency of its occurrence.

### Petite formation rate and petite viability can account for the effects of some conditions on replicative lifespan (RLS) in BY4741

In several studies, loss of mtDNA and/or MMP has been associated with reduced replicative lifespan^6,27–29^. Consistently, we observed that EtBr-pretreated mtBYtri yeast cells - 100% of which follow ΔΨ-trajectories (Fig. 3b) - are significantly shorter-lived (Fig. 4a, compare “1d LOG” and “1d LOG + EtBr”). We thus hypothesized that the survival curve of the BY4741 strain depends on the percentage of the petite cells present in the population prior to the aging experiment, and that modulation of this fraction by environmental conditions could explain their effects of on longevity. Support for this idea came from observations of the behavior of the mtBYtri under standard growth condition in nutrient-rich YPD medium. We found that replicative lifespan correlates with the time spent in logarithmic (LOG) phase prior to microfluidic analysis: prolonged growth in LOG shortens the RLS, while overnight passage through the diauxic shift increases it (Fig. 4a). RLS shortening was accompanied by an increase in the fraction of petite cells in the initial population (Fig. 4b) and the fraction of ΔΨ-lineages during aging (Fig. 4c-e). This indicates that growth in YPD promotes petite formation in mtBYtri independently of aging but this strongly affects the survival curve. Consistent with this interpretation, no significant changes in RLS were observed in the “petite-less” mtDHYtri strain under the same conditions (Fig. 4f).

**Figure 4.**
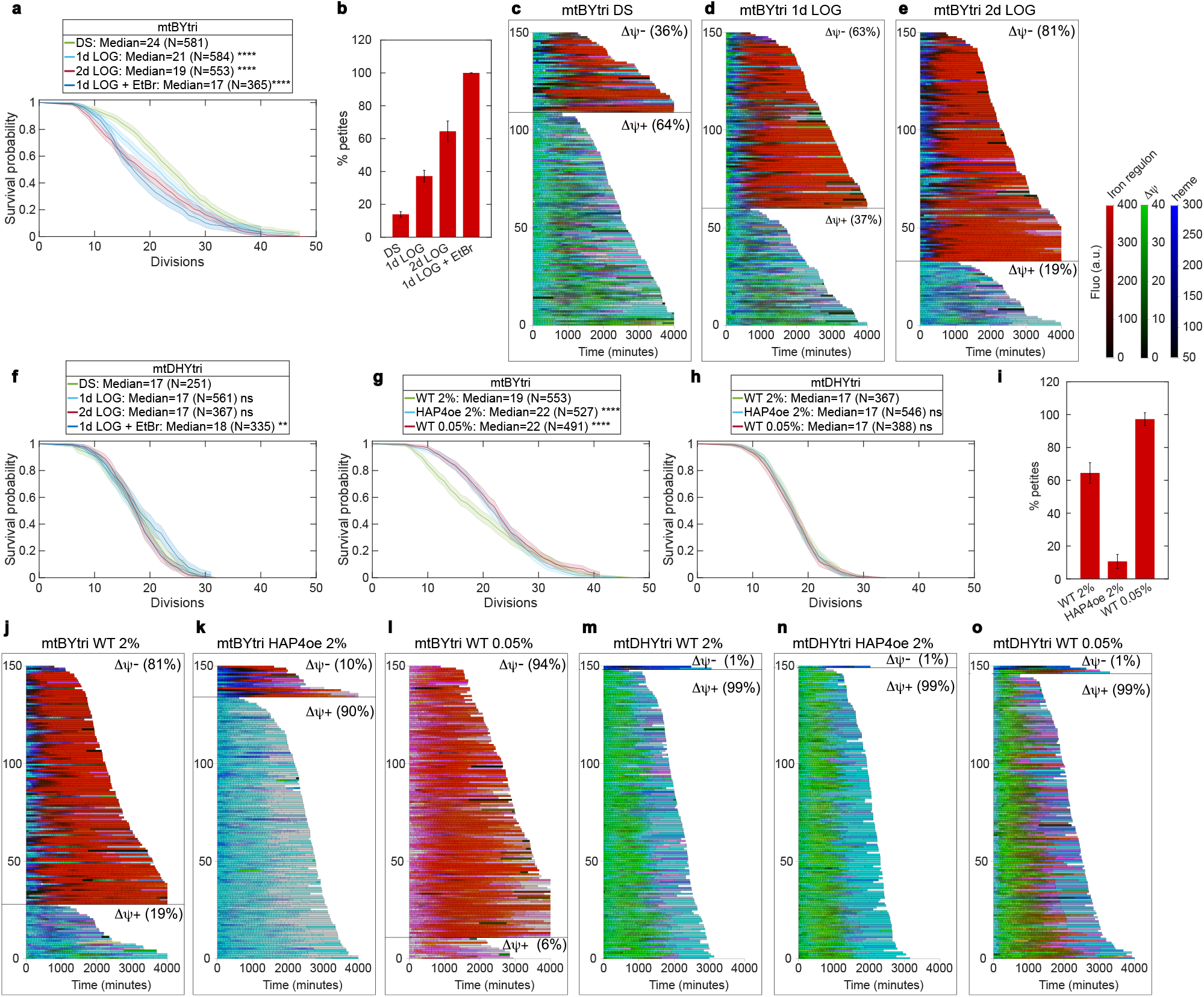
Age-independent petite formation affects longevity of the BY4741 strain. **a**, Kaplan-Meier survival curves of the indicated strains (data from at least two independent experiments). Shaded areas – 95% intervals. **** - Peto-Prentice (modified Wilcoxon) weighted log-rank test p-values < 0.0001 (1d LOG vs DS, 2d LOG vs 1d LOG, 1d LOG + EtBr vs 1d LOG). **b**, Petite frequencies of the mtBYtri strain grown in the indicated conditions. Mean +-standard deviation from two (1d LOG + EtBr), three (DS and 1d LOG) or four (2d LOG) independent measurements (total cells: DS n = 525, 1d LOG n = 657, 2d LOG n = 888, 1d LOG + EtBr n = 133). **c-e**, Single-cell trajectories (150 representative lineages randomly selected from two to three independent experiments) of the indicated strains with fluorescence data from experiments in the “mother” chip. Lineages were sorted into ΔΨ- or ΔΨ+ (numbers in parentheses indicate percentage of ΔΨ- or ΔΨ+ lineages among n lineages. DS n = 570 (3 experiments), 1d LOG n = 561 (3 experiments); 2d LOG no treatment n = 545 (3 experiments)). **f-h**, same as **a**. ns - p-values > 0.05 (mtDHYtri 1d LOG vs DS, 2d LOG vs 1d LOG, YPD vs HAP4oe, YPD vs YPD 0.05%). ** - p-values < 0.01 (mtDHYtri 1d LOG + EtBr vs 1d LOG). **** - p-values < 0.0001 (mtBYtri YPD vs HAP4oe, YPD vs YPD 0.05%). **i**, Petite frequencies of the mtBYtri strain grown in the indicated conditions for 2 days in LOG phase. Mean +-standard deviation from two (YPD 0.05%), three (HAP4oe) or four (YPD) independent measurements (total cells: YPD n = 888, HAP4oe n = 356, YPD 0.05% n = 118). **j-o**, Same as **c-e**, but with indicated strains grown for 2 days in LOG phase. 150 lineages randomly selected from one representative experiment. Numbers in parentheses indicate percentage of ΔΨ- or ΔΨ+ lineages among n lineages. YPD n = 545 (3 experiments), 1d LOG n = 561 (3 experiments); 2d LOG no treatment n = 545 (3 experiments).

Several conditions have been reported to extend replicative lifespan by reducing the proportion of aging trajectories associated with mitochondrial dysfunction and potentially improving respiration during aging. These include overexpression of the mitochondrial biogenesis factor *HAP4* and caloric restriction^6,30–33^. Consistently, we also observed that *HAP4* overexpression in the mtBYtri strain effectively abolished ΔΨ-trajectories (Fig. 4j,k) and increased median RLS (Fig. 4g). Because mtBYtri HAP4oe also exhibited a markedly reduced petite formation rate (Fig. 4i), these effects are more likely to reflect a change in the frequency with which cells enter the ΔΨ-state than a direct effect of *HAP4* on replicative aging itself. In line with this interpretation, *HAP4* overexpression had no detectable effect on RLS (Fig. 4h) or reporter dynamics (Fig. 4m,n) in the mtDHYtri strain.

By contrast, glucose restriction produced a different phenotypic outcome. Although, glucose restriction has been reported to induce mitochondrial respiration^34^, we found that after two days of logarithmic (LOG) phase growth in YPD containing 0.05% glucose, even young mtBYtri cells exhibited signs of mitochondrial dysfunction, including reduced MMP and activation of iron regulon (Fig. 4j,l, numerical data in Extended Data Fig. 5). The vast majority of these cells produced petite colonies (Fig. 4i) and aging trajectories resembling ΔΨ-cells (Fig. 4l). Despite this apparent loss of respiratory capacity, mother cells lived longer, and preCox15-NG signal in old ΔΨ-cells was higher than in cells grown in YPD with 2% glucose (Fig. 4j,l, numerical data in Extended Data Fig. 5). In addition, heme levels were increased in both young and old ΔΨ-cells under low-glucose conditions (Fig. 4j,l and Extended Data Fig. 5), further suggesting improved mitochondrial performance. These results suggest that lowering glucose concentration in rich medium promotes the viability of respiratory deficient cells both in the initial population and during aging, likely by enhancing mitochondrial performance in a respiration-independent fashion. Consistent with a petite-specific effect of glucose restriction, we observed no increase in RLS (Fig. 4h) or MMP in aged mtDHYtri cells (Fig. 4m,o). Instead, both young and aged mtDHYtri mother cells exhibited reduced MMP and activation of the iron regulon in low glucose, suggesting that caloric restriction is detrimental to mitochondrial function in respiratory-competent yeast (Fig. 4m,o and Extended Data Fig. 5). Overall, these results indicate that aging phenotype of the BY4741 strain is strongly shaped by petite cells present in the population prior to the aging experiment, in a highly condition-dependent manner.

## DISCUSSION

In this study, we uncover a major impact of an impaired mtDNA maintenance system on replicative aging in the BY4741 budding yeast strain, one of the most widely used models in longevity research. As a descendant of the S288C founder strain, BY4741 and other members of the BY series carry several mutations that compromise mitochondrial function^15,35^. Old mothers of BY strains exhibit multiple signs of mitochondrial decline, which have been interpreted as evidence for a mitochondrial aging pathway contributing to replicative senescence^1^. Our results confirm key aspects of this phenotype, including loss of MMP-dependent protein import, activation of the iron regulon, reduced heme levels, and decreased bud size in a subset of aging lineages (Fig. 1).

In contrast to previous studies, where aging cells are compared to young cells at the population level, we used a novel microfluidics-based methodological approach that enables direct comparison of aging and non-aging/rejuvenating lineages within the same experimental setting. In populations, yeast cells are subject to selection, such that the young fraction of the population is not only younger but also enriched for the fittest cells. By tracking dividing cells in the absence of selection, our approach captures slowly proliferating or arrested daughters, providing a more appropriate reference for replicatively aging mother cells.

Within this framework, we found that rejuvenating lineages lose mitochondrial function at the same rate as aging lineages and exhibit a similar signature of mitochondrial dysfunction (Fig. 2). In addition, the proportion of petite cells in the initial populations correlated with the fraction of lineages showing mitochondrial decline during aging across different conditions and genetic backgrounds (Fig. 3-4). Together, these results strongly suggest that the petite phenotype of the S288C strain and the so-called mitochondrial pathway of yeast replicative aging (“mode 2”^6^ or “round MoD”^13^ trajectories) reflect the same underlying process. This interpretation challenges the view that mitochondrial dysfunction is an intrinsic hallmark of replicative aging in yeast.

In this respect, mitochondrial dysfunction in BY4741 can be viewed as a form of clonal senescence reminiscent of telomere dysfunction. While telomere dysfunction is considered a hallmark of aging in metazoans^2^, it is generally not viewed as part of the normal replicative aging process in yeast due to efficient maintenance by telomerase^1,4,36,37^. In telomerase-deficient mutants, however, progressive telomere shortening limits both replicative lifespan and the survival of young cells^38^. Passaging of yeast cultures after complete telomerase inactivation leads to accumulation of slowly dividing/arrested cells and a gradual increase in population doubling time that is readily observed. In BY4741, by contrast, mtDNA maintenance is only partially impaired, and petite cells are counter-selected during overnight growth at saturation (Fig. 4b). As a result, growth defects are more difficult to detect at the population level in wild-type BY4741, which may explain why mitochondrial dysfunction in yeast has been considered as a feature of aging rather than as clonal senescence.

We show that prolonged propagation of mtBYtri in logarithmic phase in YPD enriches for petite cells in the population, supporting the idea that mitochondrial dysfunction arises largely independently of replicative age (Fig. 4b). Although previous studies have proposed that damaged mitochondria are preferentially retained in mother cells, thereby contributing to rejuvenation ^39–41^, our results indicate that these mechanisms are insufficient to preserve mitochondrial function over time, even in young BY4741 cells (Fig. 2).

Another striking observation of this study is that the response to mtDNA loss differs markedly between BY4741 and DHY213. Following EtBr treatment, DHY213 cells showed a milder reduction in MMP, no iron regulon activation, and preserved heme levels compared with BY4741 (Fig. 3). We further discovered that the MKT1-30G allele is largely responsible for the observed differences. Consistently, iron regulon activation is absent from previously reported transcriptomic and proteomic responses to mtDNA loss in the W303 strain^42,43^, which also carries the MKT1-30G allele. These observations suggest that the “pathological” mitochondrial dysfunction in petite BY4741 cells – characterized by strong iron dysregulation and pronounced MMP loss – is strongly influenced by the presence of the rare mkt1-30D allele and should therefore be viewed as an exception. This strain-specific feature should be taken into account when extrapolating findings from BY-derived laboratory strains to yeast more broadly or to other systems.

Mechanistically, Puf3 has been shown to promote MMP generation and proliferation of petite cells in W303 by boosting translation of mitochondrial proteins^42^. More recently, Mkt1, in complex with another RNA-binding protein Pbp1, was found to interact with Puf3 and stimulate the expression of Puf3-dependent mitochondrial proteins^24,44^. Together with these findings, our results suggest that Mkt1 (in complex with Puf3 and Pbp1) is an important component of the adaptive response to mtDNA loss, promoting MMP generation and mitochondrial function when the respiratory chain is impaired.

Intriguingly, one of the proposed human Mkt1 homologues, FAM120A^24^, shares target mRNAs with PUM1 and PUM2^45^ – human Pumilio-family homologs of yeast PUF proteins, including Puf3. Remarkably, PUM2 has been implicated in mitochondrial function during mammalian aging^46^. In addition, both PUM proteins and FAM120A were identified as potential binding partners of ATXN2 – mammalian homologue of yeast Pbp1^47^, and ATXN2 knockout in mice leads to reduced expression of multiple mitochondrial enzymes^48^. It is therefore tempting to speculate, that the role of the emerging Mkt1/Puf3/Pbp1 regulatory network in mitochondrial function could be more conserved than currently appreciated.

Despite its major role in shaping the phenotype of mitochondrial dysfunction in yeast, we found that *MKT1* does not stimulate the loss of mitochondrial function: swapping *MKT1* alleles between mtBYtri and mtDHYtri strains did not change their ΔΨ-generation frequencies (Fig. 3a,e and Fig. 3j,l). This is consistent with a previous study showing that *SAL1, CAT5* and *MIP1* determine the rate of petite formation, whereas *MKT1* only affects viability of petite cells^15^. These other alleles also likely contribute to the phenotype of ΔΨ-lineages (Fig. 3b,m), underscoring the complexity of the response to the respiratory deficiency. Thus, the apparent link between mitochondrial dysfunction and replicative aging in yeast can be profoundly influenced by background-dependent mitochondrial instability. Distinguishing intrinsic age-associated phenotypes from stochastic or genotype-dependent dysfunction will be essential for defining which alterations truly belong to the aging process.

## ACKNOWLEDGEMENTS

We thank Theo Aspert and Rama Abdulhamid for designing the geometry of the traps and creating an AutoCAD photomask for the “daughter” chip. We also thank Angela M. Chu, Joe Horecka, Joseph Schacherer, Nan Hao and Zhou Xu for sharing yeast strains and plasmids. Figure panels 1a and 2a,b were created with BioRender.com.

## AUTHOR CONTRIBUTIONS

A.M. and G.C. contributed to the conception and design of the work, analysis and interpretation of the data and manuscript preparation. A.M. performed data acquisition.

## DECLARATION OF GENERATIVE AI AND AI-ASSISTED TECHNOLOGIES IN THE WRITING PROCESS

During the preparation of this work, the authors used ChatGPT 5.1 in order to improve the clarity and fluency of the English language. After using this tool or service, the authors reviewed and edited the content and take full responsibility for the content of the publication.

## METHODS

### Strains and plasmids

Strains were developed from BY4741 (Euroscarf), DHY213 (a gift from Angela M. Chu and Joe Horecka, Ron Davis Lab) or RM11-1a (a gift from Joseph Schacherer). Gene replacements were performed by LiOAc/ssDNA/PEG transformation of the strains^49^ with DNA integration cassettes. Genetic constructs were assembled using COSPLAY protocol^50^. The detailed information on the genetic constructs are in the Dataset 1 (.dna files can be opened in SnapGene Viewer (https://www.snapgene.com/snapgene-viewer); open the “History” tab to see all the step of the creation). The tricolor versions of the strains were obtained by sequential transformations with the FIT2p-ymScarletI-LEU2 construct integrated into ChrVI:261039-261108 region, the ADH1p-SV40NLS-yiRFP-KANMX construct integrated at the ura3Δ0 locus, and RPL18Bp-preCox15-ymNeonGreen-URA3 construct integrated at the leu2Δ0 locus. BUD4 ORF was replaced with the HIS3MX6 cassette. For HAP4 overexpression, the NATMX-ADH1p construct was placed upstream of the HAP4 ORF. MKT1 allele replacements were performed using Cas9-mediated gene editing^51^. Random spores were selected using a standard protocol^52^ starting with diploid crosses between a tricolor MATa and reporter-free MATalpha strains. Allele identities were verified by Sanger sequencing.

| Name | Background | Genotype |
| --- | --- | --- |
| mtBYtri | BY4741 (S288C) | Mata *his3Δ1 leu2Δ0 met15Δ0 ura3Δ0 sal1-1 cat5-91I mip1-661A mkt1-30D hap1 ChrVIΔ(261039-261108)::FIT2p-ymScarlet-LEU2 ura3Δ0::ADH1p-NLSyiRFP-KANMX leu2Δ0::RPL18Bp-preCox15-yNeonGreen-URA3* |
| mtRMtri | RM11-1a | Mata *amn1-A1103T leu2Δ ura3Δ ho::loxP SAL1 CAT5-91M MIP1-661T MKT1-30G HAP1 ChrVIΔ(261039-261108)::FIT2p-ymScarlet-LEU2 ura3Δ0::ADH1p-NLSyiRFP-KANMX leu2Δ0::RPL18Bp-preCox15-yNeonGreen-URA3* |
| mtDHYtri | DHY213 | Mata *his3Δ1 leu2Δ0 met15Δ0 ura3Δ0 SAL1 CAT5-91M MIP1-661T MKT1-30G HAP1 RME1-INS-308A TAO3-1493Q ChrVIΔ(261039-261108)::FIT2p-ymScarlet-LEU2 ura3Δ0::ADH1p-NLSyiRFP-KANMX leu2Δ0::RPL18Bp-preCox15-yNeonGreen-URA3* |
| mtBYtri bud4Δ | BY4741 (S288C) | Mata *his3Δ1 leu2Δ0 met15Δ0 ura3Δ0 sal1-1 cat5-91I mip1-661A mkt1-30D hap1 ChrVIΔ(261039-261108)::FIT2p-ymScarlet-LEU2 ura3Δ0::ADH1p-NLSyiRFP-KANMX leu2Δ0::RPL18Bp-preCox15-yNeonGreen-URA3 bud4Δ:: HIS5MX* |
| mtBYtri<br>HAP4oe | BY4741 (S288C) | Mata his3Δ1 leu2Δ0 met15Δ0 ura3Δ0 sal1-1 cat5-91I mip1-661A mkt1-30D hap1 ChrVIΔ(261039-261108)::FIT2p-ymScarlet-LEU2 ura3Δ0::ADH1p-NLSyiRFP-KANMX leu2Δ0::RPL18Bp-preCox15-yNeonGreen-URA3 HAP4::ADH1p-HAP4 |
| mtDHYtri<br>HAP4oe | DHY213 | Mata his3Δ1 leu2Δ0 met15Δ0 ura3Δ0 SAL1 CAT5-91M MIP1-661T MKT1-30G HAP1 RME1-INS-308A TAO3-1493Q ChrVIΔ(261039-261108)::FIT2p-ymScarlet-LEU2 ura3Δ0::ADH1p-NLSyiRFP-KANMX leu2Δ0::RPL18Bp-preCox15-yNeonGreen-URA3 HAP4::ADH1p-HAP4 |
| mtBYtri<br>MKT1-30G | BY4741 (S288C) | Mata his3Δ1 leu2Δ0 met15Δ0 ura3Δ0 sal1-1 cat5-91I mip1-661A mkt1-30D hap1 ChrVIΔ(261039-261108)::FIT2p-ymScarlet-LEU2 ura3Δ0::ADH1p-NLSyiRFP-KANMX leu2Δ0::RPL18Bp-preCox15-yNeonGreen-URA3 |
| mtDHYtri<br>mkt1-30D | DHY213 | Mata his3Δ1 leu2Δ0 met15Δ0 ura3Δ0 SAL1 CAT5-91M MIP1-661T mkt1-30D HAP1 RME1-INS-308A TAO3-1493Q ChrVIΔ(261039-261108)::FIT2p-ymScarlet-LEU2 ura3Δ0::ADH1p-NLSyiRFP-KANMX leu2Δ0::RPL18Bp-preCox15-yNeonGreen-URA3 |
| BY/DHY #24 | BY4741/DHY213 | Mata his3Δ1 leu2Δ0 met15Δ0 ura3Δ0 sal1-1 CAT5-91M MIP1-661T mkt1-30D hap1 RME1-INS-308A TAO3-1493Q ChrVIΔ(261039-261108)::FIT2p-ymScarlet-LEU2 ura3Δ0::ADH1p-NLSyiRFP-KANMX leu2Δ0::RPL18Bp-preCox15-yNeonGreen-URA3 |
| BY/DHY #33 | BY4741/DHY213 | Mata his3Δ1 leu2Δ0 met15Δ0 ura3Δ0 sal1-1 cat5-91I mip1-661A mkt1-30D hap1 RME1-INS-308A TAO3-1493Q ChrVIΔ(261039-261108)::FIT2p-ymScarlet-LEU2 ura3Δ0::ADH1p-NLSyiRFP-KANMX leu2Δ0::RPL18Bp-preCox15-yNeonGreen-URA3 |
| BY/DHY #2 | BY4741/DHY213 | Matalpha his3Δ1 leu2Δ0 lys2Δ0 ura3Δ0 sal1-1 CAT5-91M MIP1-661T mkt1-30D HAP1 RME1 TAO3-1493E ChrVIΔ(261039-261108)::FIT2p-ymScarlet-LEU2 ura3Δ0::ADH1p-NLSyiRFP-KANMX leu2Δ0::RPL18Bp-preCox15-yNeonGreen-URA3 |
| BY/DHY #13 | BY4741/DHY213 | Mata his3Δ1 leu2Δ0 met15Δ0 ura3Δ0 SAL1 CAT5-91M MIP1-661T MKT1-30G HAP1 RME1 TAO3-1493E ChrVIΔ(261039-261108)::FIT2p-ymScarlet-LEU2 ura3Δ0::ADH1p-NLSyiRFP-KANMX leu2Δ0::RPL18Bp-preCox15-yNeonGreen-URA3 |

### Microfluidic chips

Microfluidic chip design and fabrication was performed as previously described^22^. “Mother chip” design was described previously^22^. Schematics of the “daughter chip” is in the Extended Data Figure 6.

### Cell culturing

For all experiments, we used YPD medium (1% yeast extract, 2%peptone, 2% or 0.05% glucose). Cells from glycerol stocks were thawed on YPD agar plates and grown for 2-3 days at 30 ºC. Typically, one or several single colonies were resuspended in YPD (2% glucose) and grown for the indicated amount of time (at least 24h) in LOG phase (never reaching OD600>2) until loading in the chip. For “DS” condition one or several single colonies were resuspended in YPD to an OD600 ∼1-2 and grown overnight (18 h), then diluted to OD600 ∼0.05 and grown 5-6h in LOG state before loading. In “EtBr pulse” experiments, 20 µL/mL ethidium bromide was added 5-6h prior to cell loading; immediately before loading EtBr-treated cells were spun down, and resuspended in 500 µL of fresh medium.

### Petite frequency analysis

For petite formation analysis, an aliquot of the cell culture was diluted and ∼100 cells were plated onto YPDGE (1% yeast extract, 2% peptone, 0.5% glucose, 3% glycerol, 2% ethanol) agar plate, grown for 4-5 days. The plates were photographed and the number of small/large colonies was counted to determine the frequency of the petite cell formation (% petite).

### Time-lapse microscopy

After baking, PDMS chip was connected to a peristaltic pump and washed with the culture medium for 1-2 hours. Then, cells were injected into the chip through the outlet by reversing the flow direction on the pump for 5-10 min, collecting cells by the barrier and changing the flow to normal. Medium flow: 5 µL/min for 1 day, then 10 µL/min for 1 another day and 15 µL/min for the rest of the experiment (3-4 days in total).

Cell imaging was performed using inverted Nikon Ti-E microscope (Spectra X Lumencor for fluorescent illumination, 60× 1.4 N.A. objective, CMOS camera Hamamatsu Orca Flash 4.0, Emission filters from the CrestOptics Spinning Disk Cicero). Brightfield images were acquired every 5 min and fluorescent images – every 20 min - for 72 h or more using Micromanager v.2.0 software. NeonGreen imaging: GFP filter wheel 50ms exposure with 4% light intensity; ScarletI imaging: RFP filter wheel 30ms exposure with 410% light intensity; iRFP imaging: FarRed filter wheel 200 ms exposure with 20% light intensity. Chip was heated at 30 ºC using custom sampler holder and objective heating devices. 4 fields of view (FOV) containing 52 “mother” traps or 6 field of views containing 27 “daughter” traps were typically collected for one condition during one experiment.

### Image processing and data analysis

For image processing we used Matlab-based DetecDiv software^22^ and other Matalb custom scripts. Some of the scripts were improved/modified using ChatGPT v.5.1, robustness of these modifications was always controlled. Using DetecDiv, we split FOVs into smaller regions of interests (ROIs) each containing a single trap. For “mother” trap ROIs we extracted cell division/RLS and mother cell segmentation information (predicted by neural networks within DetecDiv). For “daughter” trap ROIs we manually annotated cell division timing events and budding directions, then trained a SOLOv2 instance segmentation network (Matlab) to segment cells. We tracked a single cell at the bottom of one trap using a custom script. An image of an empty trap taken at the beginning of the experiment was used as a reference for background fluorescence measurements. Mean pixel intensities (background subtracted) within segmented cells were extracted and averaged across one cell cycle for plotting as trajectories (“fluorescent bars”) or swarm charts and box plots; cells which underwent less then 5 divisions and mothers that left their traps before death were excluded from the analysis. ΔΨ+/-classification was done by thresholding preCox15-NG signal: if a mother had at least two cell cycles with mean preCox15-NG fluorescence < 12 a.u. it was considered ΔΨ-, otherwise ΔΨ+. Mother cells were categorized into “large elongated” or “small round” types by visual inspection of the corresponding bud morphologies during 3-4 last divisions. For Kaplan-Meier estimates of RLS we censored clogged (to many cells within a ROI), emptied (mother left the trap before death) and still alive (neither dead nor arrested by the end of the time-lapse) cells; cells with RLS<5 or born 8h after the loading were discarded. The number of replicates, sample sizes and statistics used are detailed in figure legends. Exhaustive statistical analysis (two-sample t-test) of the data in Fig. 3g,i and Extended Data Fig. 5 are supplied as Supplementary Table 1 and 2, respectively.

