## Supplementary Figures for "Pathological mitochondrial dysfunction mimics an aging pathway in budding yeast"

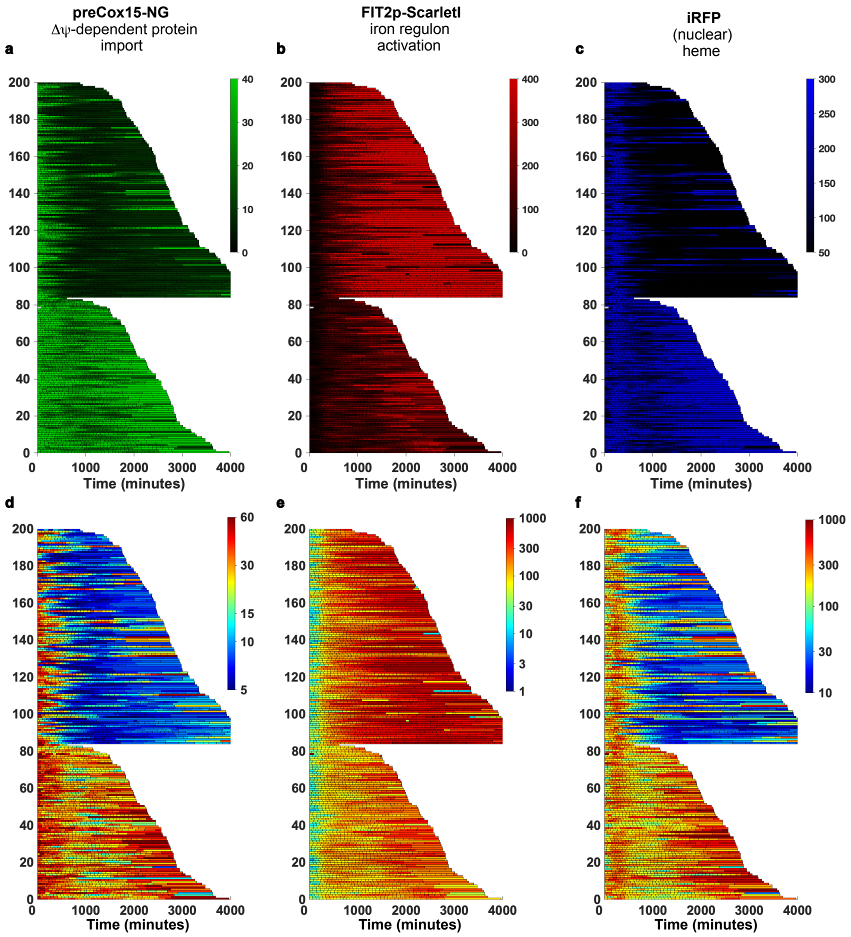


**Extended Data Figure 1.** Detailed view of the single-cell trajectories of the mtBYtri strain.

**a-f,** 200 representative mtBYtri trajectories (same as in **Fig. 1f**) only with data from one fluorescent reporter per trajectory (**a,** preCox15-Neongreen; **b,** FIT2p-ScarletI; **c,** iRFP). **d-f**, same as **a-c**, but in log10-scale and "jet" color map to accentuate smaller differences in reporter expression.

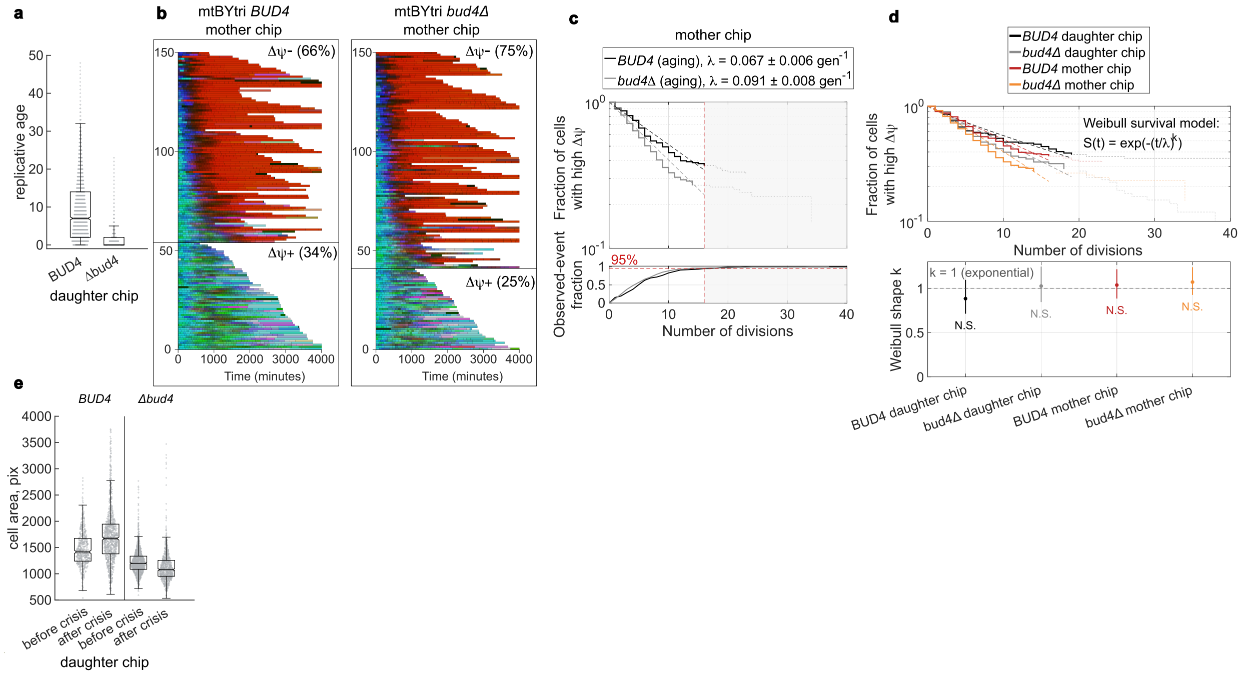

**Extended Data Figure 2.** Similarities of mitochondrial dysfunction between *BUD4* and *bud4*Δ lineages.

**a,** Replicative age during each cell cycle from the experiment in the "daughter" chip (**Fig.2e,f**) (notches on boxplots delineate area (shaded in grey) of 95% confidence intervals around the median). BUD4 median = 7, n = 2499, *bud4*Δ median = 0, n = 2045. **b,** Single-cell trajectories of the mtBYtri strains (BUD4 left, *bud4*Δ right) with fluorescence data from an experiment in the "mother" chip (150 representative randomly selected) sorted into ΔΨ- or ΔΨ+ (numbers in parentheses indicate percentage of ΔΨ- or ΔΨ+ lineages among 188 lineages from the experiment). ΔΨ+ lineages were sorted by time of yeast lifespan (in minutes); ΔΨ- lineages were sorted by time of mitochondrial dysfunction onset. 178 *bud4*Δ lineages were analyzed in the experiment. **c,** Time to ΔΨ loss in mother-chip lineages, displayed as in Fig. 2g. Kaplan-Meier estimates were computed by treating lineages that died or remained ΔΨ+ at the end of the experiment as right-censored observations. Quantitative fits were restricted to the common informative window (first 16 divisions), which contains 95% of observed events in *BUD4* mother lineages. Solid lines show the Kaplan-Meier estimates within this window; dashed lines show exponential fits; dotted extensions indicate the late censored tail that was not used for fitting. The lower panel shows the cumulative fraction of observed ΔΨ-loss events used to define the cutoff. **d**, Weibull analysis of ΔΨ-loss onset in daughter- and mother-chip lineages. Top, Kaplan-Meier survival curves for the four conditions together with Weibull fits (dashed lines), shown over the chip-specific informative windows. Bottom, Weibull shape parameter k with 95% confidence intervals for each condition; k = 1 corresponds to an exponential law with constant hazard. In all four cases, the confidence intervals overlap k = 1, indicating that the data do not reject an exponential description of ΔΨ-loss timing. **e**, Cell area (in pixels) before and after mitochondrial dysfunction onset ("before or after crisis") (notches on boxplots delineate area (shaded in grey) of 95% confidence intervals around the median). Only ΔΨ- lineages were analyzed. *BUD4* before crisis n = 478, *BUD4* after crisis n = 921, *bud4*Δ before crisis n = 1139, *bud4*Δ after crisis n = 751.

a

| Strains | Alleles | | | | |
|---|---|---|---|---|---|
| **BY4741** | sal1-1 | cat5-91I | mip1-661A | hap1 | mkt1-30D |
| **RM11-1a** | SAL1 | CAT5-91M | MIP1-661T | HAP1 | MKT1-30G |
| **DHY213** | SAL1 | CAT5-91M | MIP1-661T | HAP1 | MKT1-30G |

b

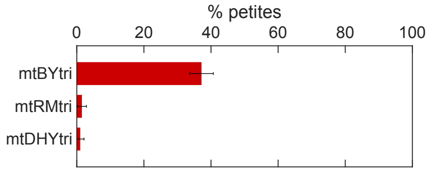


c

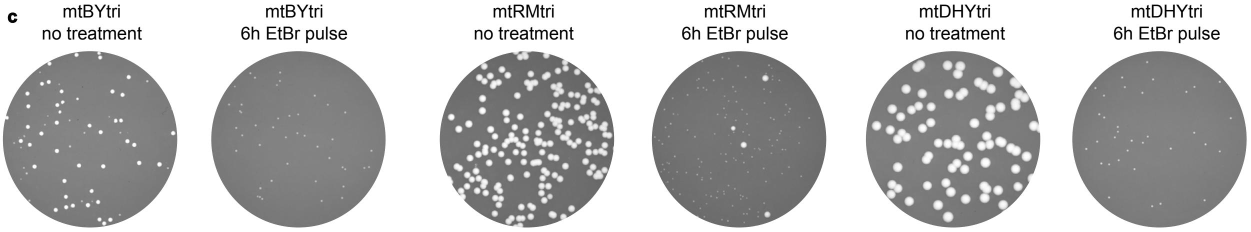


d

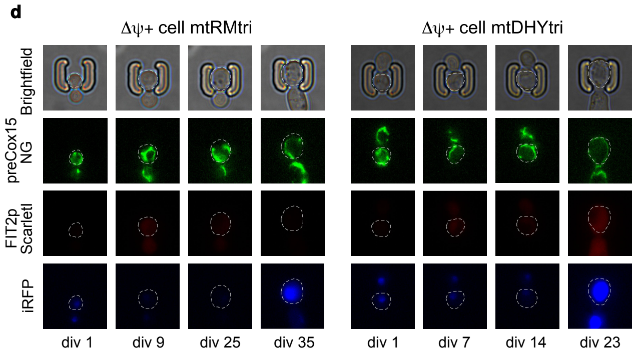


**Extended Data Figure 3. a**, Table summarizing allelic differences in the five genes affecting mitochondrial function. **b**, Petite frequencies in the indicated strains. Mean +- standard deviation from three independent measurements (total cells: mtBYtri n = 657, mtRMtri n = 465, mtDHYtri n = 717). **c,** Aliquots of cultures of the indicated strains were plated onto YPDGE and photographed after 4 days of growth. **d,** Time-lapse sequence of brightfield and fluorescent images of aging ΔΨ+ mother cell (mtRMtri and mtDHYtri as indicated). White dashed lines contour mother cells; the number of divisions is indicated below.

**a**

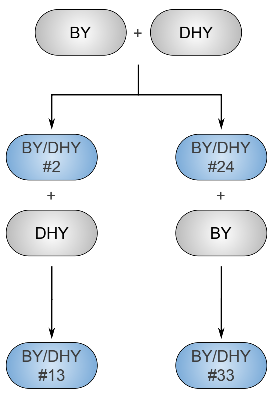


**b**

| Strain | CAT5 | MIP1 | SAL1 | MKT1 | HAP1 | RME1 | TAO3 | iron regulon activation in Δψ− | heme levels in Δψ− | Δψ− trajectories |
|---|---|---|---|---|---|---|---|---|---|---|
| BY4741 | BY | BY | BY | BY | BY | BY | BY | + | - | |
| DHY213 | DHY | DHY | DHY | DHY | DHY | DHY | DHY | - | + | |
| BY/DHY #24 | DHY | DHY | BY | BY | BY | DHY | DHY | + | - | |
| BY/DHY #33 | BY | BY | BY | BY | BY | DHY | DHY | + | - | |
| BY/DHY #2 | DHY | DHY | BY | BY | DHY | BY | BY | + | - | |
| BY/DHY #13 | DHY | DHY | DHY | DHY | DHY | BY | BY | - | + | |

**c**

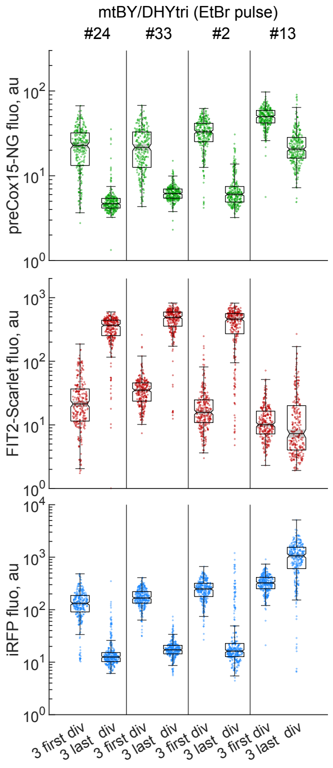


**Extended Data Figure 4.** Contribution of different BY/DHY alleles to changes in MMP, iron regulon activation and heme levels upon loss of respiratory function.

**a**, Schematic demonstrating the process of generation of crosses ##2, 13, 24 and 33 between BY and DHY strains. **b**, Table summarizing the allelic differences between the indicated strains (BY – allele from the BY4741 strain, DHY - allele from the DHY213 strain). The "ΔΨ- trajectories" column contains single-cell trajectories with fluorescence data (50 representative lineages) of the indicated strains from one experiment in the "mother" chip after EtBr 6h pulse.). **c,** Log10-transformed fluorescence (in a.u.) during three first and three last divisions (notches on boxplots delineate area (shaded in grey) of 95% confidence intervals around the median). mtBY/DHYtri #24 n = 282, #33 n = 282, #2 n = 279, #13 n = 282 (1 experiment).

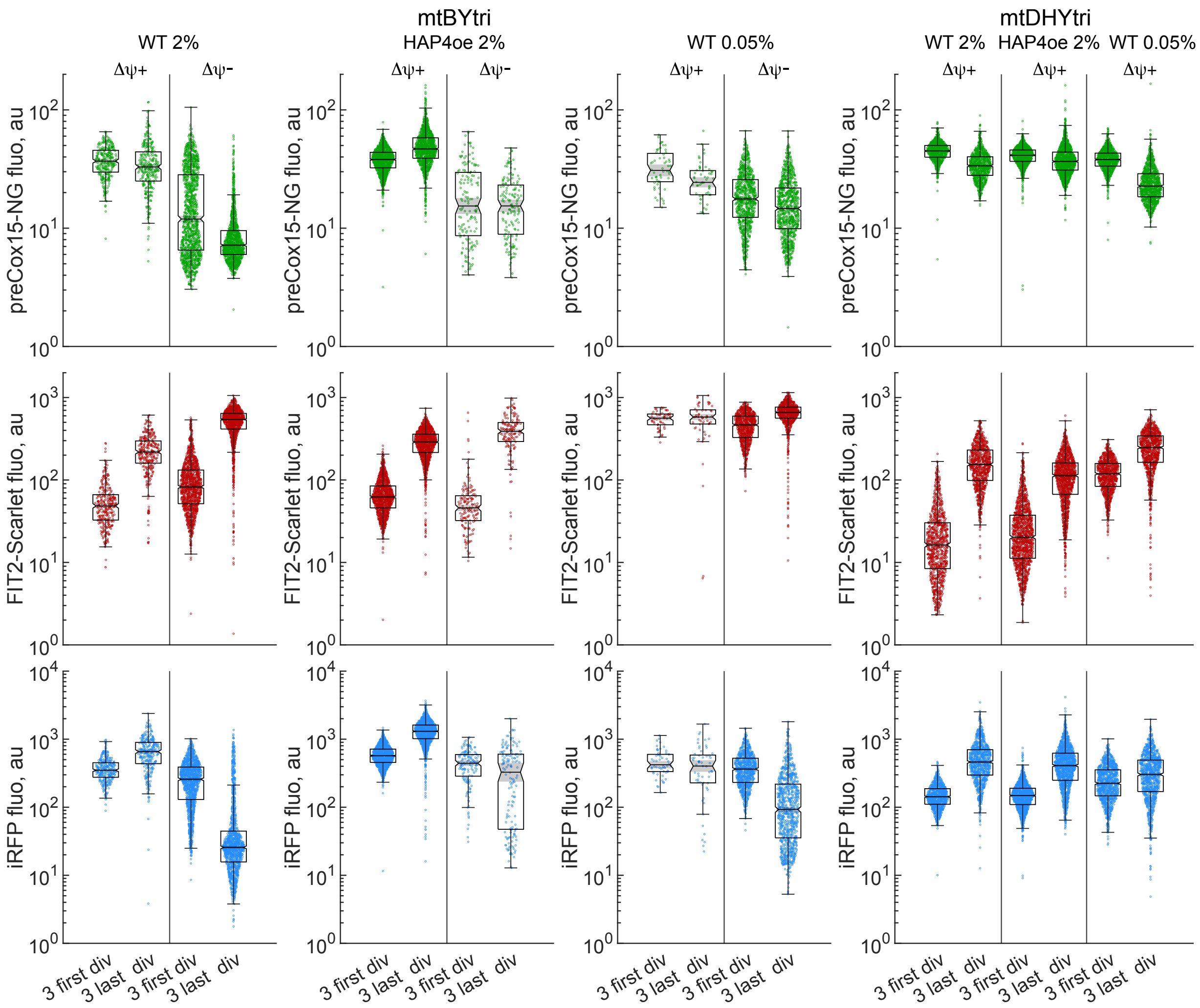


**Extended Data Figure 5.** Numerical data for the Figure 4

Log10-transformed fluorescence (in a.u.) during three first and three last divisions (notches on boxplots delineate area (shaded in grey) of 95% confidence intervals around the median). mtBYtri YPD ΔΨ+ n = 315, ΔΨ- n = 1320 (3 experiments); mtBYtri HAP4oe ΔΨ+ n = 1359, ΔΨ- n = 156 (3 experiments); mtBYtri YPD 0.05% ΔΨ+ n =57, ΔΨ- n =939 (2 experiments); mtDHYtri YPD ΔΨ+ n = 948 (2 experiments); mtDHYtri HAP4oe ΔΨ+ n = 1443 (3 experiments); mtDHYtri YPD 0.05% ΔΨ+ n = 1056 (2 experiments). Statistical analysis (two-sample t-test) of the data are in Supplementary Table 2.

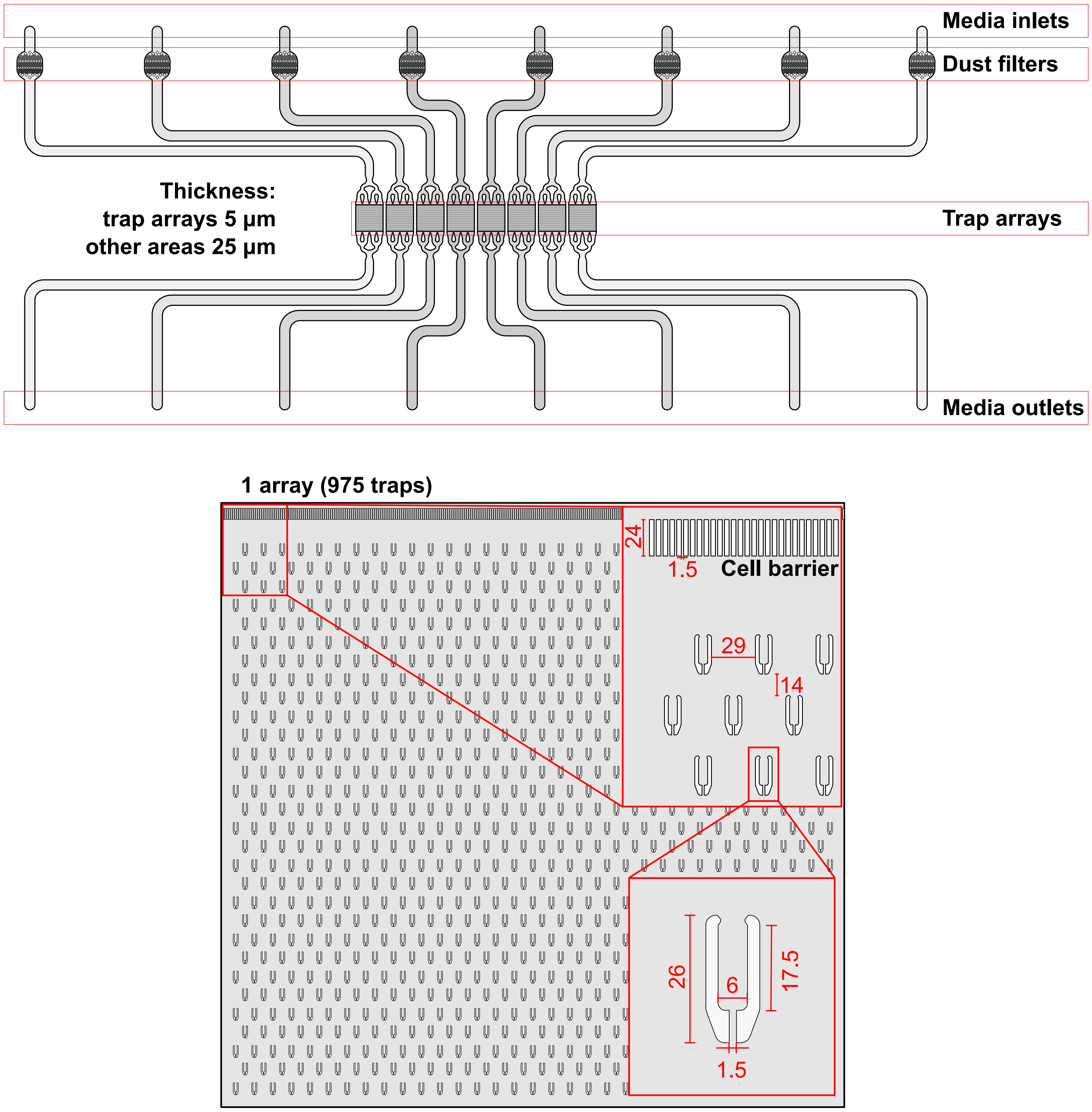


**Extended Data Figure 6.** Schematics of the "daughter" microfluidic device. Dimensions are in µm.
